# MADS-box factor AGL16 negatively regulates drought resistance via stomatal density and stomatal movement

**DOI:** 10.1101/723106

**Authors:** Ping-Xia Zhao, Zi-Qing Miao, Jing Zhang, Qian-Qian Liu, Cheng-Bin Xiang

**Affiliations:** School of Life Sciences and Division of Molecular & Cell Biophysics, Hefei National Science Center for Physical Sciences at the Microscale, University of Science and Technology of China, The Innovation Academy of Seed Design, Chinese Academy of Sciences, Hefei, Anhui Province 230027, China

**Author notes:** Corresponding author: Cheng-Bin Xiang, School of Life Sciences, University of Science and Technology of China, Hefei, Anhui 230027, China.

**Keywords:** AGL16, drought resistance, stomatal density, ABA, MADS-box

## Abstract

Drought is one of the most severe environmental factors limiting plant growth and productivity. Plants respond to drought by closing stomata to reduce water loss. The molecular mechanisms underlying plant drought resistance are very complex and yet to be fully understood. While much research attention has been focused on the positive regulation of stomatal closure, less is known about its negative regulation, equally important in this reversible process. Here we show that the MADS-box transcriptional factor AGL16 acts as a negative regulator in drought resistance by regulating both stomatal density and movement. Loss-of-function mutant *agl16* was more resistant to drought stress with higher relative water content, which was attributed to a reduced leaf stomatal density and more sensitive stomatal closure due to a higher leaf ABA level compared with wild type, while *AGL16* overexpression lines displayed the opposite phenotypes. *AGL16* is preferentially expressed in guard cells and down regulated in response to drought stress. The expression of *CYP707A3* and *AAO3* in ABA metabolism and *SDD1* in stomatal development was altered by AGL16 as shown in *agl16* and overexpression lines. Chromatin immunoprecipitation, transient transactivation, and yeast-one-hybrid assays demonstrated that AGL16 bound the CArG motif in the promoter of the *CYP707A3*, *AAO3*, and *SDD1* to regulate their transcription, and therefore alter leaf stomatal density and ABA level. Taken together, AGL16 acts as a negative regulator of drought resistance by modulating leaf stomatal density and ABA accumulation.

## INTRODUCTION

Drought is the most adverse abiotic stress that directly threatens crop growth and productivity (Basu et al., 2016; Le and McQueen-Mason, 2006; Zhu, 2016). Drought stress disturbs normal physiological and morphological traits from cellular metabolism to changes in growth rates in plants by altering water potential and cell turgor (Anjum et al., 2011; Vurukonda et al., 2016). It is crucial to understand the molecular mechanism and adaptation by which crops cope with water deficit, and take actions to improve plant resistance against environmental pressures for food security of the world (Anjum et al., 2011; Tester and Langridge, 2010).

ABA is the key phytohormone governing plant drought response. It used to be considered that ABA is synthesized in roots and transported to leaves to induce stomatal closure (Wilkinson and Davies, 2010). More studies suggest that ABA can be synthesized in specific cells of vascular tissues and exported to guard cells, or autonomously synthesized in guard cells and ABA biosynthesis is upregulated by positive feedback in response to ABA (Bauer et al., 2013; Kuromori et al., 2010; Kuromori et al., 2014). Upon water deficit, ABA accumulation is increased in guard cells, which triggered stomatal closure resulting in the decline of transpiration rate of leaves. ABA biosynthesis and catabolism are thought to be dominant factors determining endogenous ABA levels in plants which is directly related to plant tolerance (Lim et al., 2015; Nambara and Marion-Poll, 2005). Most genes involved in ABA biosynthesis such as *ABA1*, *ABA2*, *ABA3*, *NCED* and *AAO3* are induced by abiotic stresses. 9-cis-epoxycarotenoid dioxygenase (NCED) is considered to be a key enzyme catalyzing the rate-limiting step in ABA biosynthesis (Iuchi et al., 2001; Wan and Li, 2006). *NCED* was upregulated in guard cells after exposure to ABA (Bauer et al., 2013), but it is unknown whether a complete ABA biosynthesis pathway is present in those cells (Iuchi et al., 2001; Malcheska et al., 2017). *ABA2* encodes a short-chain dehydrogenase/reductase-like enzyme that converts xanthoxin to ABA-aldehyde, which is finally converted to ABA by abscisic aldehyde oxidase3 (*AAO3*) that is activated by molybdenum cofactor (MoCo) sulfurase ABA3 (Nambara and Marion-Poll, 2005; Schwartz et al., 2003). The ABA content is significantly decreased in Arabidopsis *aba2* and *aao3* mutants, which show enhanced sensitivity to drought (Nambara et al., 1998; Seo et al., 2000; Wang et al., 2018). Recently, the essential nutrient sulfate has been confirmed to induce stomatal closure (Batool et al., 2018; Chen et al., 2019), probably by affecting ABA biosynthesis via cysteine as sulfur donor for MoCo modification by ABA3.

*CYP707A* encodes the key cytochrome P450 enzyme ABA 8′-hydroxylase (ABA 8′-OH) in ABA catabolism that modulates cellular ABA levels. There are four *CYP707A* genes in the subfamily, *CYP707A1*-*CYP707A4*. *CYP707A3* is most responsive to dehydration and rehydration, and loss-of-function mutant *cyp707a3* exhibits a higher ABA level and enhanced tolerance to drought (Umezawa et al., 2006). *CYP707A1* and *CYP707A2* play crucial roles in modulating endogenous ABA levels during seed dormancy and germination in Arabidopsis (Matilla et al., 2015; Millar et al., 2006; Okamoto et al., 2006). Moreover, *CYP707A1* was also shown to be involved in post germination growth and stomatal conductance that was remarkably reduced in cyp*707a1* mutant compared to that of wild type (Okamoto et al., 2009; Zhu et al., 2011).

Stomata, formed by two specialized epidermal guard cells, are the main channels regulating CO_2_ influx from external environmental and H_2_O loss from leaves (Kim et al., 2010). During stomatal differentiation, bHLH proteins have been identified as major positive regulators of stomatal lineage fate. SPEECHLESS (SPCH), MUTE, FAMA determine the initial transition of a protodermal cell to an meristmoid mother cell, the meristemoid to the guard mother cell, and direct the symmetric division to form guard cells (MacAlister et al., 2007; Ohashi-Ito and Bergmann, 2006; Pillitteri and Torii, 2007). FOUR LIPS (FLP) and MYB88 negatively regulate the division from GMC to GCs. Division and differentiation were more severely affected, resulting an extension of GMC division in *flp myb88* double mutants (Lai et al., 2005; Xie et al., 2010). STOMATAL DENSITY AND DISTRIBUTION 1 (*SDD1*) encodes a subtilisin-like serine protease which likely processes propeptides into ligands to activate the TMM-ER complex that regulates the MAPK cascade (Berger and Altmann, 2000; Von Groll et al., 2002; Yoo et al., 2010). It appears to act independently of the EPF peptides to regulate stomatal development (Franks et al., 2015b; Hunt et al., 2010; Hunt and Gray, 2009). Compared to the wild type, stomatal density on the abaxial leaf surface was about 2.5-fold higher in *sdd1* mutant, and overexpression of *SDD1* led to a 2 to 3-fold decrease in stomatal density (SchluEter et al., 2003; Von Groll et al., 2002). Overexpression of maize *SDD1* (*ZmSDD1*) reduced stomatal density resulting the decrease of stomatal conductance and transpiration rate, and exhibiting an enhanced drought resistance in maize (Liu et al., 2015). GTL1 is a positive regulator that modulates stomatal density by transrepression of SDD1 to regulate WUE and drought tolerance (Yoo et al., 2010).

AGAMOUS-LIKE16 (AGL16) transcription factor is a member of MIKC-type MADS-box family in Arabidopsis. Recently, it was reported that AGL16 was targeted by miR824 in the regulation of satellite meristemoid lineage of stomatal development. The transgenic plants expressing miR824-resistant AGL16 mRNA increased the incidence of stomata in higher-order complexes (Kutter et al., 2007). Furthermore, miR824-regulated AGL16 contributes to flowering time repression in a long-day photoperiod (Hu et al., 2014).

In this study, we continue to unravel the molecular mechanisms of EDT1/HDG11-conferred drought resistance (Yu et al., 2008) and focus on the EDT1-downregulated MADS-box factor AGL16. We demonstrated that transcription factor AGL16 acted as a negative regulator of drought resistance by transcriptionally repressing *AAO3* and *SDD1* and activating *CYP707A3* to regulate leaf stomatal density and ABA level.

## RESULTS

### *AGL16* negatively regulates drought resistance

To investigate the function of AGL16, we obtained a T-DNA insertion mutant (Salk_104701, *agl16*) from Arabidopsis Biological Resource Center (ABRC) and generated *AGL16* overexpression lines (OX) in wild type background as well as functional complementation lines (FC) in *agl16*, and their expression level of *AGL16* was confirmed (Supplemental Figure S1).

Plants grown in soil did not show any obvious morphological difference among *agl16*, wild type (Col-0), OX and FC lines under normal growth conditions. For drought assay, 4-week-old soil-grown plants were withheld watering for 17 days before rewatering to recover. The *agl16* mutant displayed drought resistant phenotype compared with wild type, whereas OX lines were more sensitive to drought, and FC lines showed similar phenotype to Col-0 (Figure 1A). The mutant *agl16* exhibited a higher survival rate after rehydration recovery, while the OX line had a lower survival rate compared with wild type (Figure 1B).

**Figure 1.**
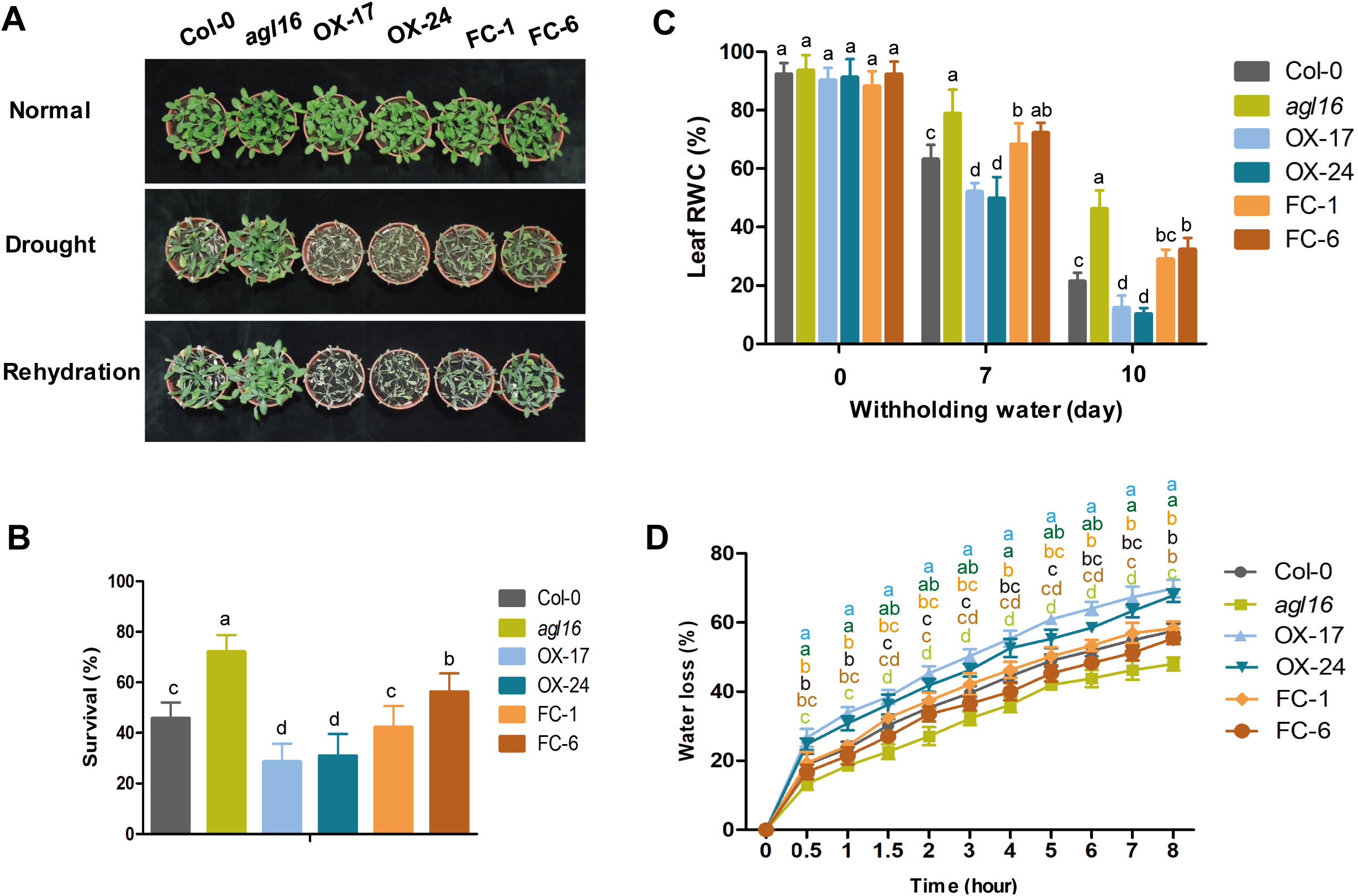
AGL16 negatively regulates drought resistance in Arabidopsis. (A-B) Drought resistance assays in soil. 4-week-old Col-0, *agl16*, OX and FC lines were grown under normal condition or with drought treatment for 14 days, then rewatered. Photographs were taken at the end of drought treatment and after rehydration (A) and survival rate was calculated (B). Values are mean ± SD (n=3 replicates, 80 plants/replicate). Different letters indicate a significant difference by Student’s t-test (P < 0.05) (C) Relative water content of leaves. 4-week-old soil-grown seedlings were subjected to drought stress by withholding water for indicated days and leaf fresh weight was measured and leaf RWC (%) were calculated. Values are mean ± SD (n=3 replicates, 30 plants per genotype). Different letters indicate a significant difference by Student’s t-test (P < 0.05). (D) Water loss rate of detached leaves. Detached rosette leaves were dried under 22□ and 60% relative humidity and weighed at the indicated time points and water loss rate (%) was calculated. Values are mean ± SD (n=3 replicates, 10 plants per genotype). Different letters indicate a significant difference by Student’s t-test (P < 0.05).

To estimate plants transpiration under drought stress, we measured leaf relative water content (RWC). *agl16* mutant held a higher RWC than Col-0 after withholding water for 7 or 10 days, whereas OX lines showed much lower RWC (Figure 1C). We also measured water loss rate of detached rosette leaves. Again, *agl16* mutant showed a lower rate of water loss than Col-0 at different time points, while the OX lines had higher rates (Figure 1D). Furthermore, soil-grown *agl16* plants displayed higher leaf temperature than Col-0, whereas OX lines showed lower leaf temperature, consistent with their water loss rate (Supplemental Figure S2). These results of different genotypes correlate well with their phenotype to drought, demonstrating that AGL16 negatively regulates drought resistance in Arabidopsis.

### AGL16 regulates stomatal density and stomatal movement

To further investigate the mechanism by which AGL16 regulates drought resistance, we examined stomatal density and stomatal movement, two major contributors of drought resistance. We found that *agl16* mutant had a reduced stomatal density compared to Col-0, while the OX-17 and OX-24 lines displayed an increased stomatal density (Figure 2A-B). Nevertheless, compared with Col-0, the density of pavement cells exhibited no remarkable changes in *agl16* mutant as well as in OX-17, OX-24 lines (Figure 2C). In addition, stomatal index showed a significant difference in *agl16* mutant and OX-24 compared to Col-0 by statistical analyses (Figure 2D).

**Figure 2.**
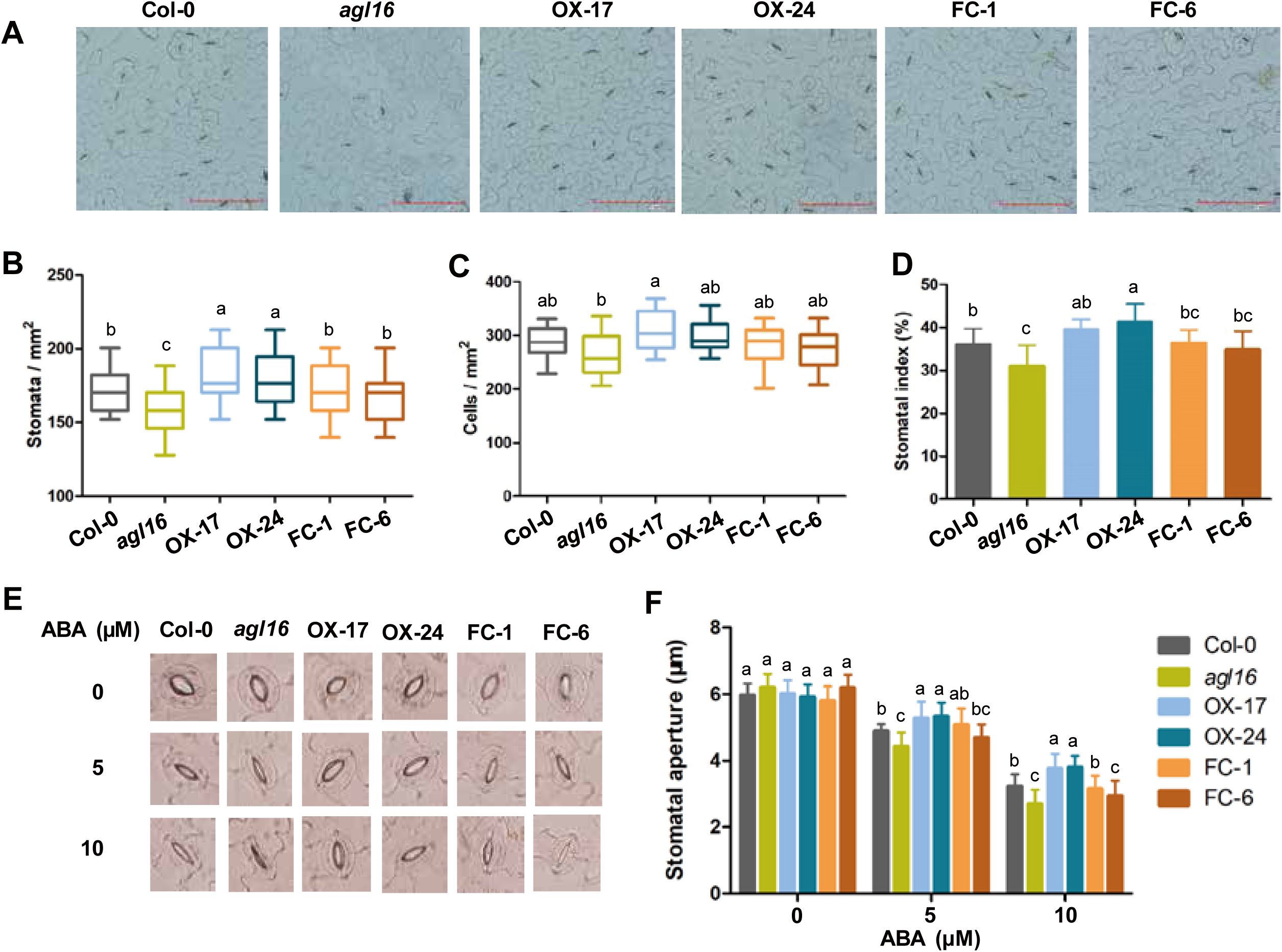
AGL16 regulates stomatal density and stomatal movement. (A-D) AGL16 affects stomatal density and stomatal index. Stomatal number of rosette leaves with the same leaf position was observed by microscope (A), and stomatal density (B), epidermal cell density (C) and stomatal index (D) were measured. Values are mean ± SD (n=20 plants per genotype, 5 images/plants). Different letters indicate significant difference by Student’s t-test (P < 0.05). (E-F) Stomatal aperture response to ABA. Rosette leaves with the same leaf position were soaked in stomatal opening buffer for 4 hours, then transferred to buffer with 5, 10 µM ABA for 2 hours before photographs were taken (E). Stomatal aperture was measured under microscope (F). Values are mean ± SD (n=20 plants per genotype, 5 images/plants). Different letters indicate a significant difference by Student’s t-test (P < 0.05).

We also examined the sensitivity of guard cell to ABA. The width of stomatal aperture was measured after 0, 5 or 10 µM ABA treatment. Under normal condition, stomatal aperture of Col-0, *agl16*, OX and FC lines were roughly the same. In response to 5 or 10 µM ABA treatment, *agl16* showed more sensitive stomatal closure than Col-0, whereas OX lines were less sensitive to ABA, and FC lines had no significant difference compared to Col-0 (Figure 2E-F). Together, these results demonstrate that AGL16 regulates drought resistance via both stomatal density and stomatal movement.

### AGL16 affects ABA content and ROS activity under drought stress

Considering that ABA plays a central role in drought stress signaling and that the sensitivity of stomata closure was altered by AGL16, we suspected that ABA content might be affected. Therefore, we measured leaf ABA content in Col-0, *agl16*, OX, and FC lines. The ABA levels did not show a significant difference among the genotypes in the absence of stress treatment. When withheld watering for 10 days, a significant increase in ABA content of all genotypes were observed. However, *agl16* mutant accumulated a higher ABA content than Col-0, while OX lines had significantly lower ABA content (Figure 3A). These results suggest that the elevated ABA content contributes to drought resistance of *agl16* mutant.

**Figure 3.**
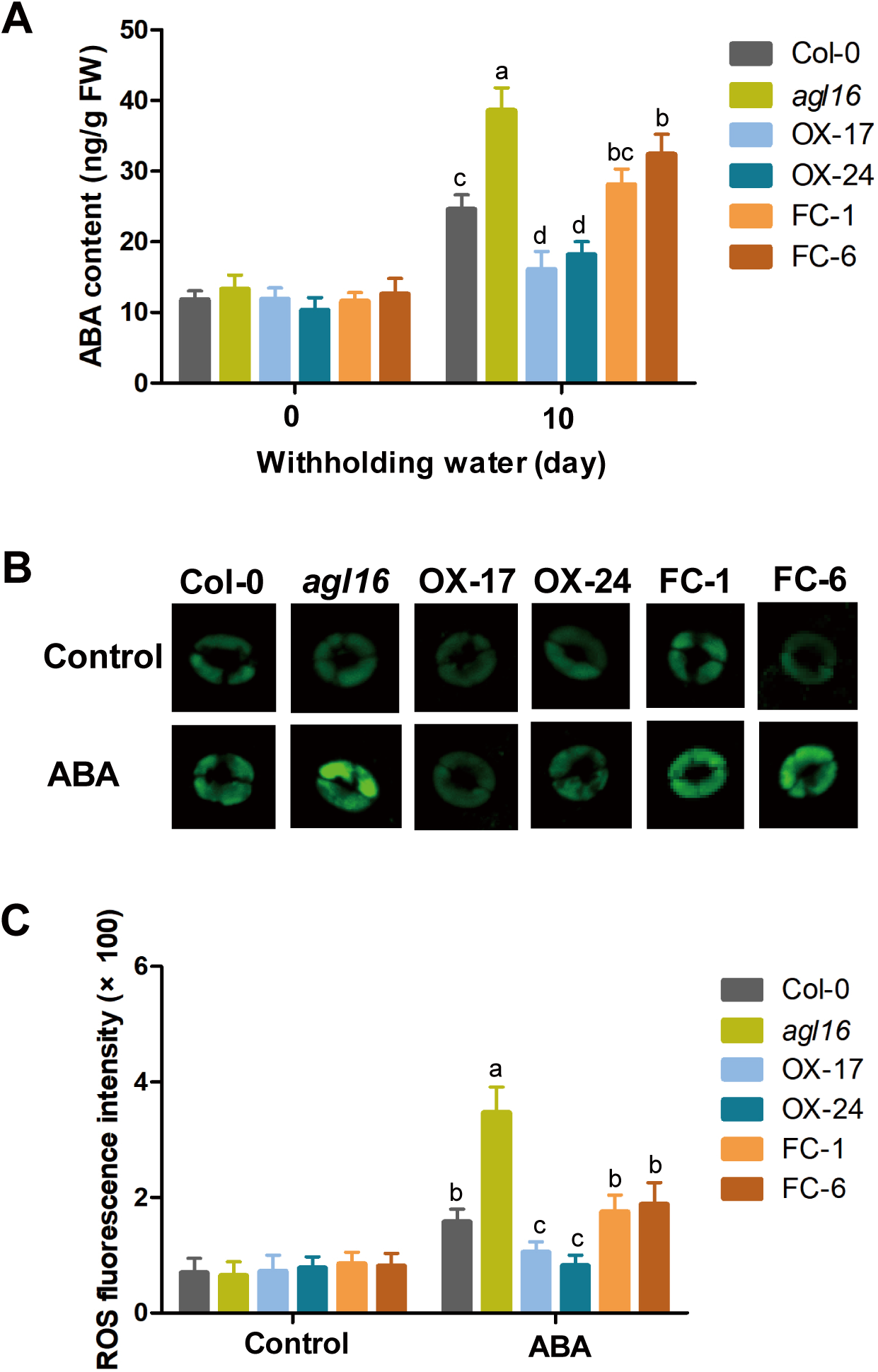
AGL16 affects ABA content and ROS activity under drought stress. (A) Leaf ABA content of Col-0, *agl16*, OX and FC lines. 4-week old plants were grown in soil under normal condition or withholding water for 10 days, then ABA content was detected. Values are mean ± SD (n=3 replicates, 20 plants per genotype). Different letters indicate significant difference by Student’s t-test (P < 0.05). (B-C) Guard cell ROS activity. Epidermal stripes from 4-week-old soil-grown plants of Col-0, *agl16*, OX and FC lines were incubated in fluorescent labeling solution with H_2_DCF-DA plus 0 or 10 μM ABA for 1 hour at 37°C and in the dark before photographs were taken (B). Bar=100 μm. Fluorescent intensity was quantified with ImageJ software (NIH, USA) (C). ROS level in guard cells was quantified based on fluorescent intensity. The fluorescence intensities in Col-0, *agl16*, OX and FC lines were multiplied by 100, respectively. Values are mean ± SD (n=3 replicates, 20 plants per genotype, 5 images/plants). Different letters indicate significant difference by Student’s t-test (P < 0.05).

ABA is known to induce the production of reactive oxygen species (ROS) in guard cells. To determine whether AGL16 affects ROS production, we examined ROS activity in guard cells. Under normal condition, all the lines did not show significantly altered ROS production in guard cells. When exposed to exogenous ABA, ROS-specific fluorescence of guard cells was significantly increased in *agl16* mutant, decreased in OX lines, and similar in FC lines compared to Col-0 (Figure 3B-C), consistent with that an elevated ABA level triggers ROS production.

### AGL16 is preferentially expressed in stomata and localized in nucleus

To analyze the tissue expression pattern of *AGL16*, we examined the transcript level of *AGL16* in different tissues of 4-week-old Arabidopsis plants by quantitative RT-PCR. Higher expression levels of *AGL16* were observed in leaves, stem, and siliques than that in other tissues (Figure 4A). To analyze the expression pattern in details, we generated *AGL16pro:*GUS transgenic plants with a 2kb *AGL16* promoter to drive β-glucuronidase (GUS) reporter gene and found that the *AGL16* gene was preferentially expressed in guard cells in the leaf abaxial epidermis by GUS staining (Figure 4B), consistent with its role in guard cells.

**Figure 4.**
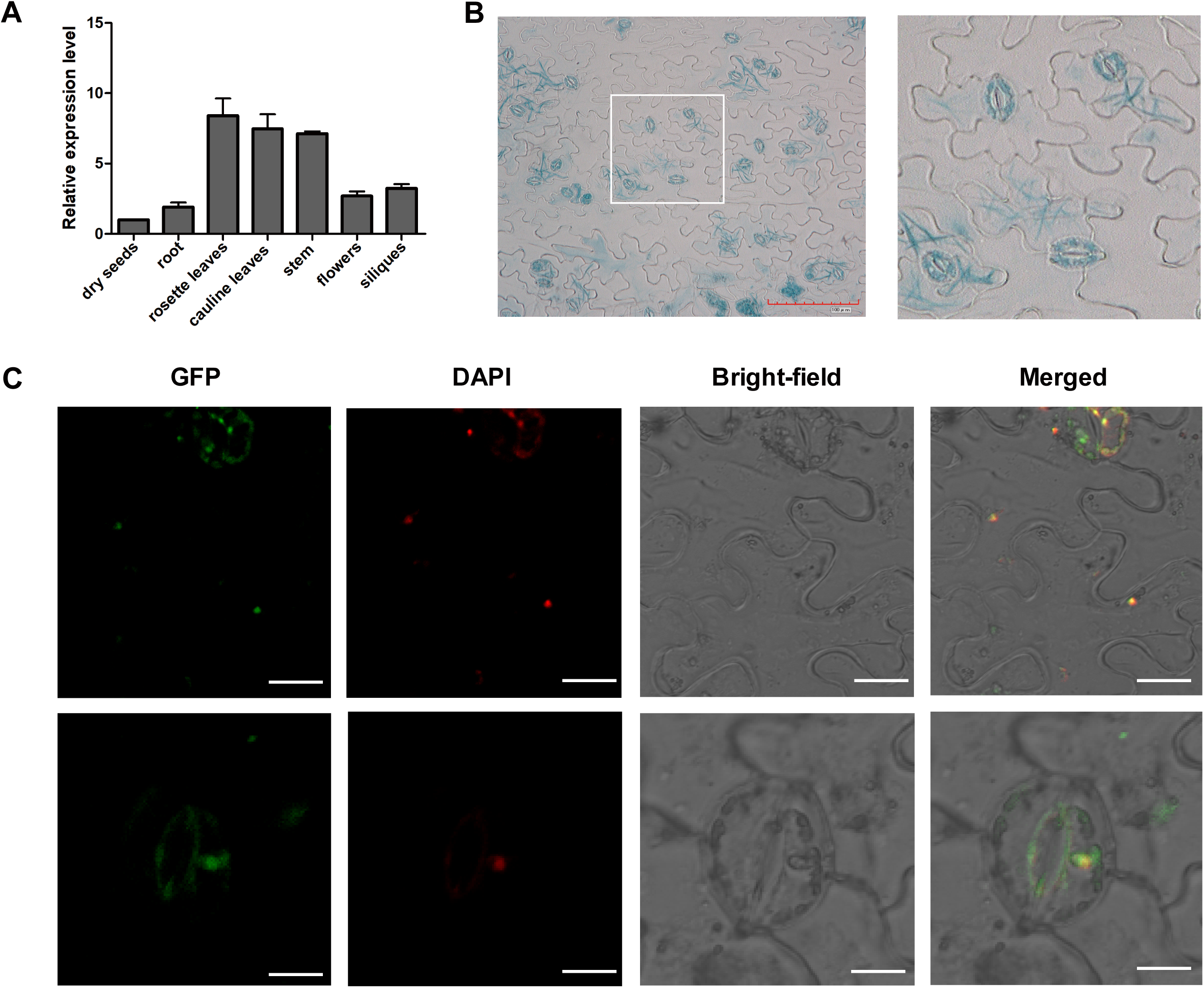
AGL16 is preferentially expressed in guard cell and localized in nucleus. (A) Tissue expression pattern of *AGL16.* Different tissues of 4-week-old adult plants were used to isolate RNA and detect the transcription level of *AGL16* by quantitative RT-PCR analysis. Values are mean ± SD (n=3 replicates) (B) GUS activity in the abaxial epidermis of *AGL16pro: GUS* transgenic plants. The boxed area was enlarged and shown on the right. Bar=100 μm (C) Subcellular localization of AGL16. *35S:AGL16-GFP* fusion proteins transiently expressed in *Nicotiana benthamiana* epidermal cells, then kept in darkness at 28°C for 48 h, and fluorescence was observed under confocal laser scanning microscope. Bar=25 μm.

To investigate the subcellular localization, we constructed *35S:AGL16-GFP* vector that was transiently expressed in *Nicotiana benthamiana* epidermal cells. We found that AGL16 protein was localized to nuclei in guard cells and pavement cells by confocal laser scanning microscope (Figure 4C).

Furthermore, we investigated the response of *AGL16* to drought stress and ABA. The transcript level of *AGL16* was gradually downregulated in response to PEG and ABA treatment (Supplemental Figure S3), but slowly increased by ABA treatment after 12h (Supplemental Figure S3B).

### AGL16 affects the expression of genes in ABA catabolic and biosynthetic pathway

To understand how AGL16 affect ABA content, we examined the expression of genes in ABA catabolic and biosynthetic pathway. The transcript levels of the ABA catabolic genes *CYP707A1*, *CYP707A2*, *CYP707A4* were not markedly affected in Col-0, *agl16*, OX lines (Figure 5). However, the expression of *CYP707A3* was significantly upregulated in the OX line and downregulated in *agl16* under both control and drought treatment. Drought treatment significantly elevated *CYP707A3* transcript level compared with control. In addition, we found that the expression of *AAO3* in ABA biosynthesis pathway was markedly elevated in *agl16* mutant but decreased in OX line. These results indicate that AGL16 may directly target the genes of ABA catabolic and biosynthetic pathway and therefore affect ABA content.

**Figure 5.**
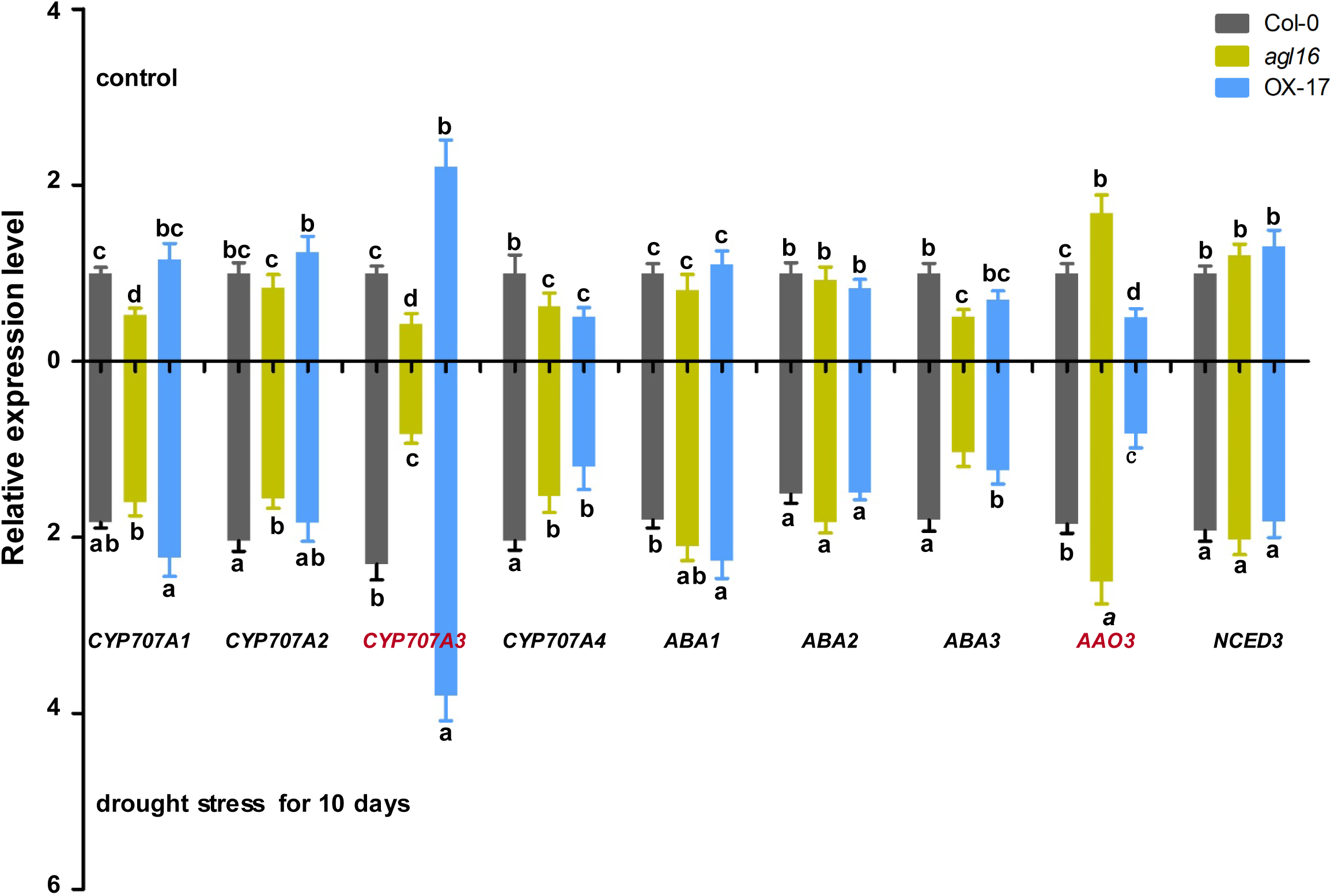
The expression of *CYP707A3* and *AAO3* is altered by AGL16. Transcript level of ABA catabolic pathway genes *CYP707A1*, *CYP707A2*, *CYP707A3*, *CYP707A4* and ABA biosynthetic pathway genes *ABA1*, *ABA2*, *ABA3*, *AAO3*, *NCED3* were measured in 4-week-old Col-0, *agl16*, and OX plants being withheld water for 0 or 10 days by quantitative RT-PCR analysis. Values are mean ± SD (n=3 replicates). Different letters indicate a significant difference by Student’s t-test (P < 0.05).

Furthermore, quantitative RT-PCR analyses showed that the transcript levels of ABA-responsive genes *ABI1*, *ABF1*, *RD29B*, and *RD22* were upregulated in *agl16* mutant, and downregulated in OX line under drought stress, which were not seen in the absence of drought (Supplemental Figure S4). This result suggests that AGL16 also negatively modulates ABA responsiveness possibly through altered ABA level.

We also analyzed the expression of downstream genes involved in stomatal movement including K^+^ channels *GORK*, *KAT1*, *KAT2*, *AKT1*, *AKT2*, and anion channel *SLAC1* by quantitative RT-PCR. When exposed to drought, the expression levels of *KAT1*, *KAT2*, *AKT1*, *AKT2* were not significantly altered. Nonetheless, the transcript abundance of *GORK* was significantly increased in *agl16* mutant compared with Col-0 (Supplemental Figure S6).

### AGL16 affects the expression of *SDD1* involved in stomatal development

We have demonstrated that AGL16 affects stomatal density. To explore the underlying molecular mechanism, we detected the transcript levels of stomatal development-related genes including *SDD1*, *ERECTA*, *TMM*, *YODA*, *MPK3*, *FAMA* in Col-0, *agl16*, OX-17 lines. The transcript abundance of *SDD1* in the shoots of 7-day-old seedling was much higher in *agl16* mutant compared with Col-0, and decreased in OX line (Figure 6A). Similarly, the expression of YODA, MPK3, FAMA was obviously elevated in *agl16* mutant, and slightly reduced in OX line (Figure 6D-F). These results indicate that AGL16 likely affects stomatal density by regulating the expression of *SDD1*, a known factor to alter stomatal density.

**Figure 6.**
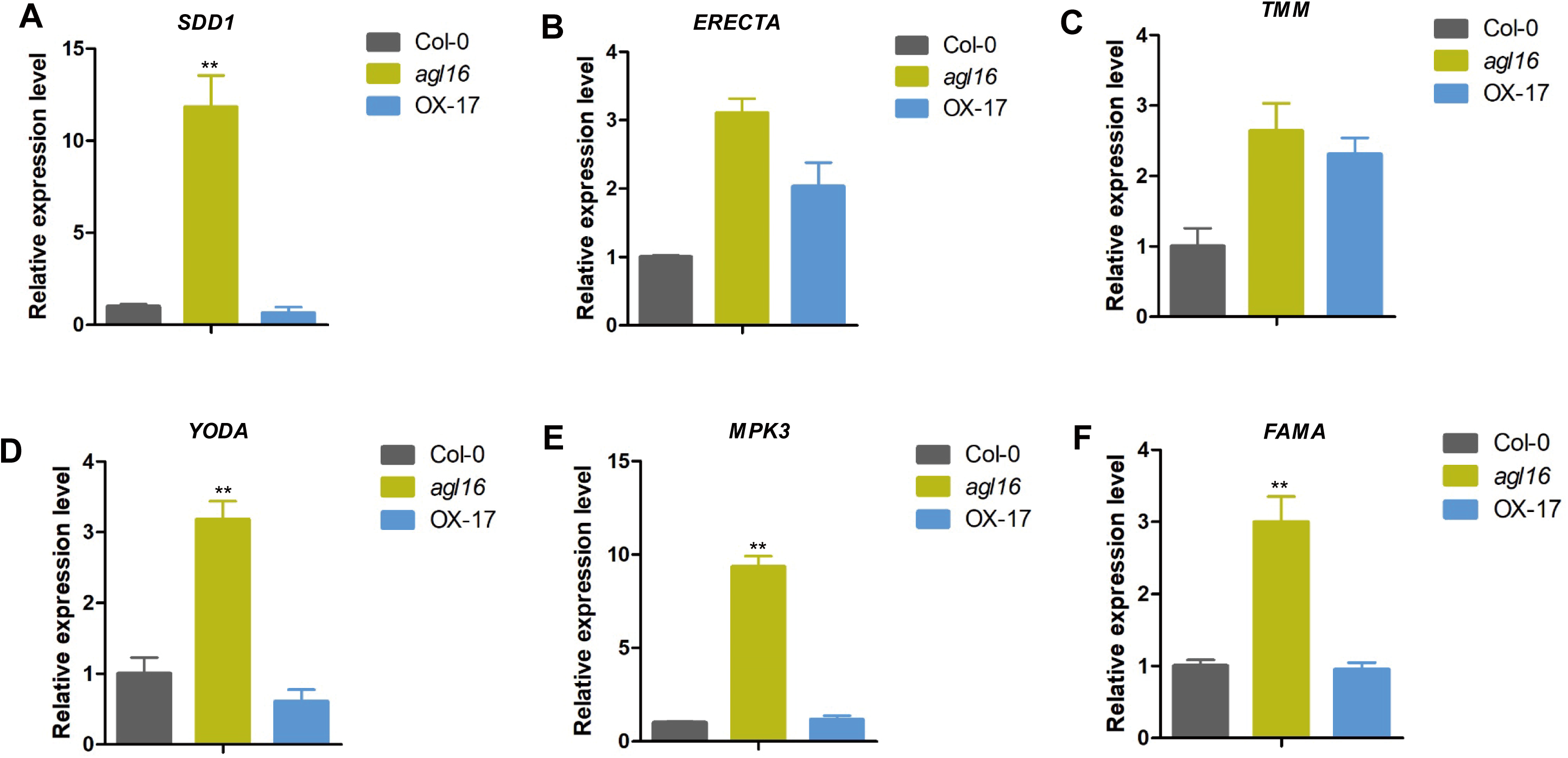
AGL16 affects the expression of stomatal development genes. Transcript level of stomatal development genes *SDD1* (A), *ERECTA* (B), *TMM* (C), *YODA* (D), *MPK3* (E), *FAMA* (F) were detected in 7-day-old Col-0, *agl16*, OX seedlings by quantitative RT-PCR analysis, respectively. Values are mean ± SD (n=3 replicates). **P < 0.01.

### AGL16 transcriptionally regulates *CYP707A3, AAO3* and *SDD1*

As a transcription factor, AGL16 may bind to the promoter of its target genes and thus directly regulate their expression. We found two putative AGL16-binding sites (CArG motifs) in each 1.6 kb promoter regions of *CYP707A3* and *AAO3*, and one putative site in the promoter of *SDD1* (Figure 7–9A). To investigate whether *CYP707A3*, *AAO3* and *SDD1* were directly regulated by AGL16, we performed chromatin immunoprecipitation (ChIP) assays. About 200 bp promoter fragments contained CArG motif of *CYP707A3*, *AAO3* and *SDD1* were markedly enriched in the *35S:HA-AGL16* transgenic plants, but not in Col-0 by PCR and quantitative PCR (Figure 7–9B). To confirm this, we further conducted transient transactivation assays in Arabidopsis mesophyll protoplasts. The coding sequence of AGL16 was driven by the cauliflower mosaic virus (CaMV) 35S promoter as an effector. The 1600 bp promoter of *CYP707A3*, *AAO3* and *SDD1* were respectively fused to the luciferase reporter gene (Figure 7–9C). The expression of the *CYP707A3* promoter-driven luciferase (LUC) was activated when both effector and reporter were transformed into Arabidopsis mesophyll protoplasts. Meanwhile, AGL16 was able to suppress the expression of the *SDD1* promoter-driven luciferase (LUC) reporter in Arabidopsis protoplasts as shown in Figure 9D. These results indicate that AGL16 can bind to the promoter of *CYP707A3*, *AAO3* and *SDD1* to regulate their expression *in vivo*.

**Figure 7.**
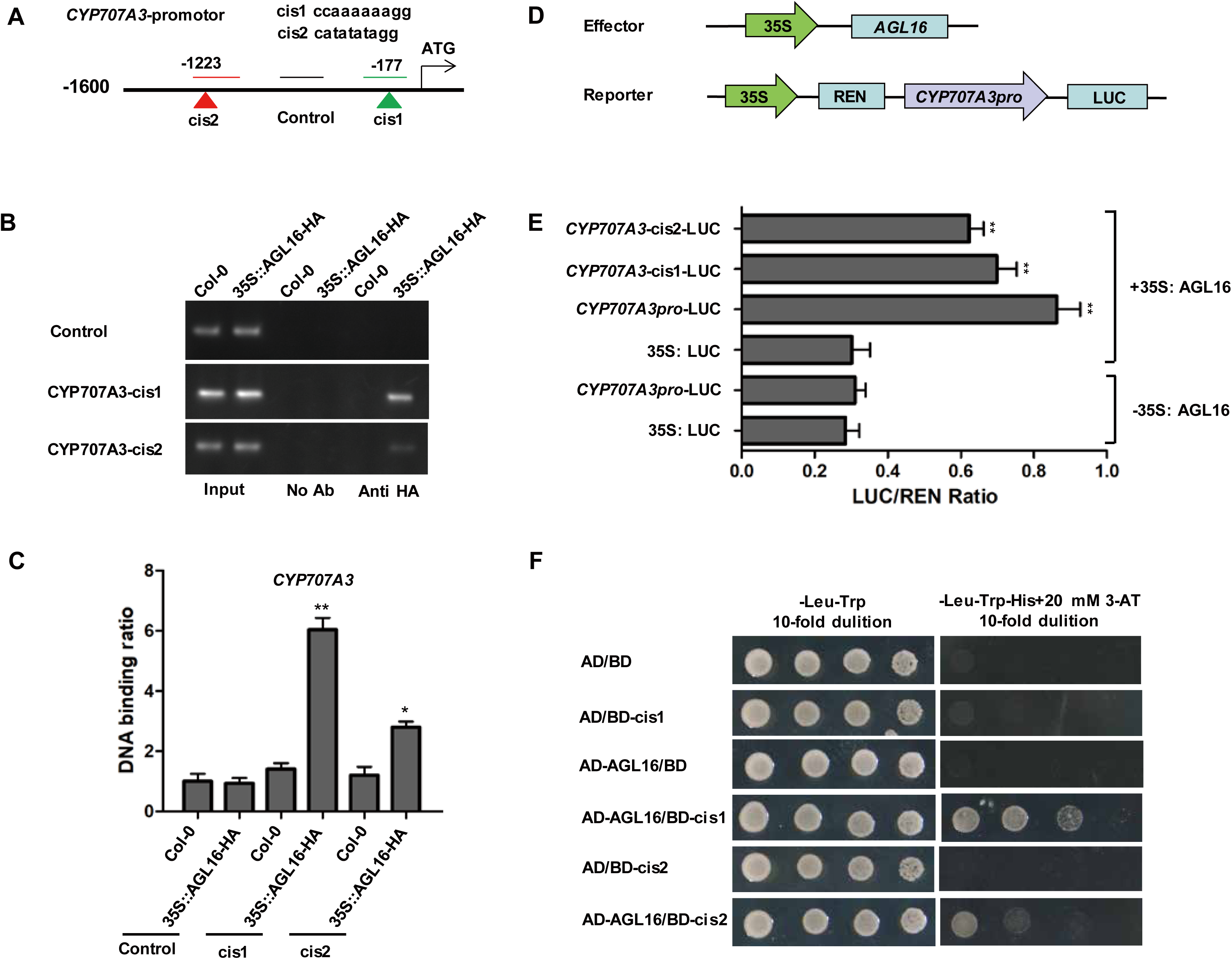
AGL16 binds and activates *CYP707A3* promoter *in vivo*. (A) Schematic representation of *CYP707A3* promoter with candidate AGL16 binding sites. AGL16 binding sites are indicated with green and red triangles in *CYP707A3* promoter. Number represents the precise location of AGL16 binding sites. Short lines mark the regions amplified in PCR and quantitative RT-PCR analysis, while the black line marks the control region without AGL16 binding site. (B-C) ChIP-PCR assay. 4-week-old soil-grown plants of Col-0 and *35S: HA-AGL16* were used for the ChIP assay. A region of *CYP707A3* that does not contain AGL16 binding site was used as a control. The enrichment was detected by PCR (B) and quantitative RT-PCR (C). Values are mean ± SD (n=3 replicates, Student’s t-test P values, *P<0.05, **P<0.01). (D-E) Transient transactivation assays. The coding sequence of AGL16 was driven by the CaMV 35S promoter as an effector. *CYP707A3* promoter and the 26 bp binding cis-element from the promoter were respectively fused to the luciferase gene as reporters (D). Both effector and reporter were transfected into Arabidopsis mesophyll protoplasts. The activities of firefly LUC and REN luciferase were measured by a dual-luciferase reporter assays, and LUC/REN ratio was calculated. pGreenII 62-SK (−35S: AGL16) and pGreenII 0800-LUC (35S: LUC) were transformed into mesophyll protoplasts as negative controls (these were not shown in the figure). The activity of REN Luciferase serves as an internal control. Values are mean ± SD (n=3 replicates, Student’s t-test P values, *P<0.05, **P<0.01). (F) Yeast-one-hybrid assay. The coding sequence of AGL16 was cloned into pGADT7 vector (AD) and the binding sites of *CYP707A3* promoter were cloned into pHIS2 (BD) vector. These constructs were transformed into Y187 yeast strain, and observed their growth on SD/-Trp-Leu medium with or without His plus 20 mM 3-AT. The combination of AD/BD, AD-AGL16/BD, AD/BD-cis1, AD/BD-cis2 were used as negative controls.

**Figure 8.**
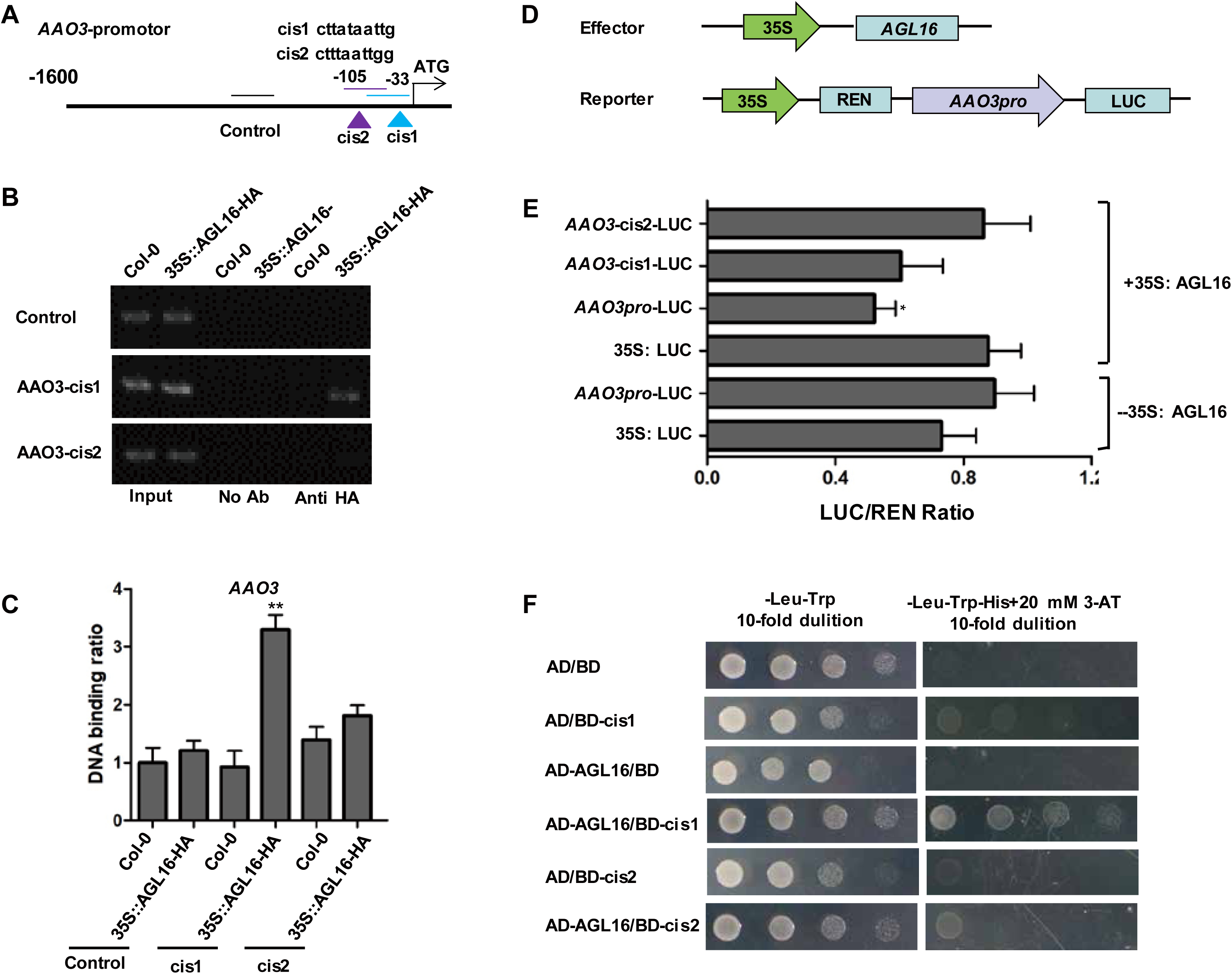
AGL16 binds and represses *AAO3* promoter *in vivo*. (A) Schematic representation of *AAO3* promoter with candidate AGL16 binding sites. AGL16 binding site is indicated with blue and purple triangle in *AAO3* promoter. Number represents the precise location of AGL16 binding sites. Short lines mark the regions amplified in PCR and quantitative RT-PCR analysis, while the black line marks the control region without AGL16 binding site. (B-C) ChIP-PCR assay. 7-day-old Col-0 and *35S: HA-AGL16* seedlings were used for the ChIP assay. A region of *AAO3* that does not contain AGL16 binding site was used as a control. The enrichment was detected by PCR (B) and quantitative RT-PCR (C). Values are mean ± SD (n=3 replicates, Student’s t-test P values, *P<0.05, **P<0.01). (D-E) Transient transactivation assays. The coding sequence of AGL16 was driven by the CaMV 35S promoter as an effector. *AAO3* promoter and the 26 bp binding cis-element from the promoter were respectively fused to the luciferase gene as reporters. Both effector and reporter were transfected into Arabidopsis mesophyll protoplasts. The activities of firefly LUC and REN luciferase were measured by a dual-luciferase reporter assays, and LUC/REN ratio was calculated. pGreenII 62-SK (−35S: AGL16) and pGreenII 0800-LUC (35S: LUC) were transformed into mesophyll protoplasts as negative controls. The activity of REN Luciferase is an internal control. Values are mean ± SD (n=3 replicates, Student’s t-test P values, *P<0.05, **P<0.01). (F) Yeast-one-hybrid assay. The coding sequence of AGL16 was cloned into pGADT7 vector (AD) and the binding site of *AAO3* promoter was cloned into pHIS2 (BD) vector. These constructs were transformed into Y187 yeast strain, and observed their growth on SD/-Trp-Leu medium with or without His plus 20 mM 3-AT. The combination of AD/BD, AD-AGL16/BD, AD/BD-cis1, AD/BD-cis2 were used as negative controls.

**Figure 9.**
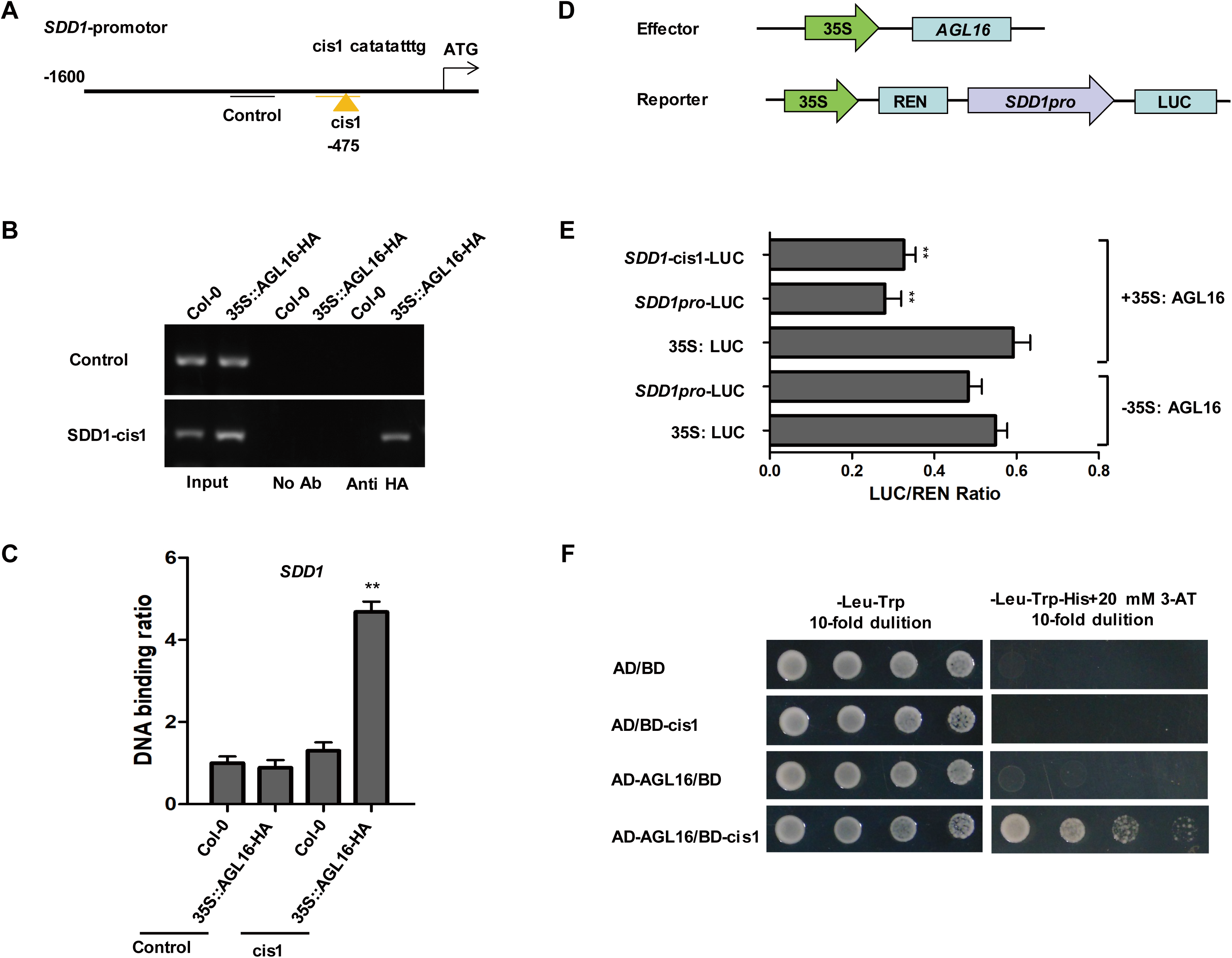
AGL16 binds and represses *SDD1* promoter *in vivo*. (A) Schematic representation of *SDD1* promoter with candidate AGL16 binding sites. AGL16 binding site is indicated with orange triangle in *SDD1* promoter. Number represents the precise location of AGL16 binding sites; Short lines mark the regions amplified in PCR and quantitative RT-PCR analysis, while the black line marks the control region without AGL16 binding site. (B-C) ChIP-PCR assay. 7-day-old Col-0 and *35S: HA-AGL16* seedlings were used for the ChIP assay. A region of *SDD1* that does not contain AGL16 binding site was used as a control. The enrichment was detected by PCR (B) and quantitative RT-PCR (C). Values are mean ± SD (n=3 replicates, Student’s t-test P values, *P<0.05, **P<0.01). (D-E) Transient transactivation assays. The coding sequence of AGL16 was driven by the CaMV 35S promoter as an effector. *SDD1* promoter and the 26 bp binding cis-element from the promoter were respectively fused to the luciferase gene as reporters. Both effector and reporter were transfected into Arabidopsis mesophyll protoplasts. The activities of firefly LUC and REN luciferase were examined by a dual-luciferase reporter assays, and LUC/REN ratio was calculated. pGreenII 62-SK (−35S: AGL16) and pGreenII 0800-LUC (35S: LUC) were transformed into mesophyll protoplasts as a negative controls. The activity of REN Luciferase is an internal control. Values are mean ± SD (n=3 replicates, Student’s t-test P values, *P<0.05, **P<0.01). (F) Yeast-one-hybrid assay. The coding sequence of AGL16 was cloned into pGADT7 vector (AD) and the binding site of *SDD1* promoter was cloned into pHIS2 (BD) vector. These constructs were transformed into Y187 yeast strain, and observed their growth on SD/-Trp-Leu medium with or without His plus 20 mM 3-AT. The combination of AD/BD, AD-AGL16/BD, AD/BD-cis1 were used as negative controls.

Furthermore, we performed yeast-one-hybrid (Y1H) assays to confirm the binding in yeast. As expected, Y1H results show that AGL16 can interact with two CArG-binding motifs of *CYP707A3* promoter, and one site of *AAO3* or *SDD1* promoter respectively (Figure 7–9E). Together, the above results suggest that AGL16 enhances the expression of *CYP707A3* and represses the expression of *SDD1* and *AAO3* by binding to the CArG motifs in their promoters.

## DISCUSSION

While positive regulation of drought resistance has drawn much of the research attention, less is known about the negative regulation. In this study, we have demonstrated that AGL16 acts as a negative regulator in drought resistance by modulating leaf stomatal density and ABA level.

Reducing stomatal density decreases transpiration and improves water use efficiency, which has been demonstrated by Arabidopsis *edt1* (Yu et al., 2008) and epidermal patterning factor mutants (Doheny-Adams et al., 2012; Franks et al., 2015a) as well as by transgenic crops with reduced stomatal density (Hughes et al., 2017; Yin et al., 2017; Yu et al., 2013; Yu et al., 2016; Zheng et al., 2017).

Stomatal density is determined mostly by changes in leaf epidermal cell size (Carins Murphy et al., 2016) or stomatal development. The reduced stomatal density of *edt1* is caused by enlarged leaf cells due to enhanced endoreduplication (Guo et al., 2019). Our study showed that stomatal index of *agl16* was slightly changed (Figure 2D), indicating that AGL16 regulates stomatal density largely through stomatal development rather than epidermal pavement cell growth, which coincided with *sdd1* mutant that increased the stomatal index and disoriented spacing divisions, but is distinct from *tmm*, *yoda*, and *er*-family mutants in stomatal phenotype (Berger and Altmann, 2000; Bergmann, 2004; Bergmann and Sack, 2007). These results are also consistent with the role of SDD1 in the regulation of asymmetric divisions to produce stomatal meristoids (Berger and Altmann, 2000; Von Groll et al., 2002). Moreover, miR824 regulates *AGL16* and affects satellite meristemoid lineage of stomatal development (Kutter et al., 2007). Our data show that the transcript abundance of *SDD1*, *YODA*, *MPK3* is elevated in *agl16* mutant, and decreased in OX line (Figures 5A, D, E), suggesting that AGL16 functions upstream of these regulatory factors and is consistent with the negative regulatory pathway in stomatal development (Bergmann and Sack, 2007).

When exposed to drought stress, plants respond to increase ABA level to rapidly mediate stomatal closure to prevent water loss (Kim et al., 2010). In this study, we revealed that AGL16 acts as a negative regulator of ABA accumulation (Figure 3A), which further affects the expression of ABA signal pathway genes including *ABI1* and *ABF1* (Supplemental Figure 5), consequently leading to the downstream events of stomatal closure including the efflux of Ca^2+^, the activation of S-type (SLAC1), R-type anion channels and GORK channel that lead to the efflux of Cl^−^, malate^2−^, NO_3_^−^ and K^+^ (Daszkowska-Golec and Szarejko, 2013; Hosy et al., 2003; Vahisalu et al., 2008) as well as H_2_O_2_ production in guard cells (Kim et al., 2010; Pei et al., 2000; Wang and Song, 2008).

Stomatal movement is a reversible process. Stomata close upon drought stress and open when drought stress is gone. This dynamic process must be finely modulated in order to balance between growth and drought response. Therefore, negative regulation of drought response is equally important. Arabidopsis ERF7 and MYB60 were reported as negative regulators of drought resistance by controlling stomatal closure (Oh et al., 2011; Song et al., 2005). A rice zinc finger transcription factor was shown to negatively regulate stomatal closure by modulating genes for H_2_O_2_ homeostasis (Huang et al., 2009). A recent report shows that the transcription factor HAT1 negatively regulates ABA synthesis and signaling in Arabidopsis in response to drought (Tan et al., 2018). AGL16 contributes one more means for plants to fine tune stomatal movement.

Our data show that AGL16 acts as a transcriptional repressor for *SDD1* and *AAO3* or activator for *CYP707A3.* It is not uncommon for a transcriptional factor to be both activator and repressor depending on its target promoter context. For instance, WRKY53 can act either as transcriptional activator or repressor depending on the promoter sequences that it interacts with (Miao et al., 2004). WRKY6 was shown to act as an activator on *PR1* expression during plant senescence and pathogen defense, and it also suppresses its own and *WRKY42* expression (Miao et al., 2004; Robatzek and Somssich, 2001; Robatzek and Somssich, 2002). To fulfill this double role, AGL16 is likely assisted by other nuclear protein partners.

Taken together, our study shows that AGL16 functions as a new negative regulator of drought resistance via stomatal density and ABA accumulation. A working model is proposed for AGL16 (Figure10), where AGL16 enhances the expression of *CYP707A3* and represses the expression of *AAO3*, *SDD1* by binding to the CArG motifs in their promoters, and consequently changes ABA accumulation by affecting both ABA degradation and biosynthesis. On the other hand, AGL16 may regulate stomatal density in early leaf development in response to environmental changes. Collectively, AGL16 negatively regulate drought resistance by altering stomatal density and leaf ABA accumulation, serving as an important modulator balancing between drought resistance and growth. Moreover, AGL16 should provide a novel candidate for modifying crops via readily available genome editing tools.

**Figure 10.**
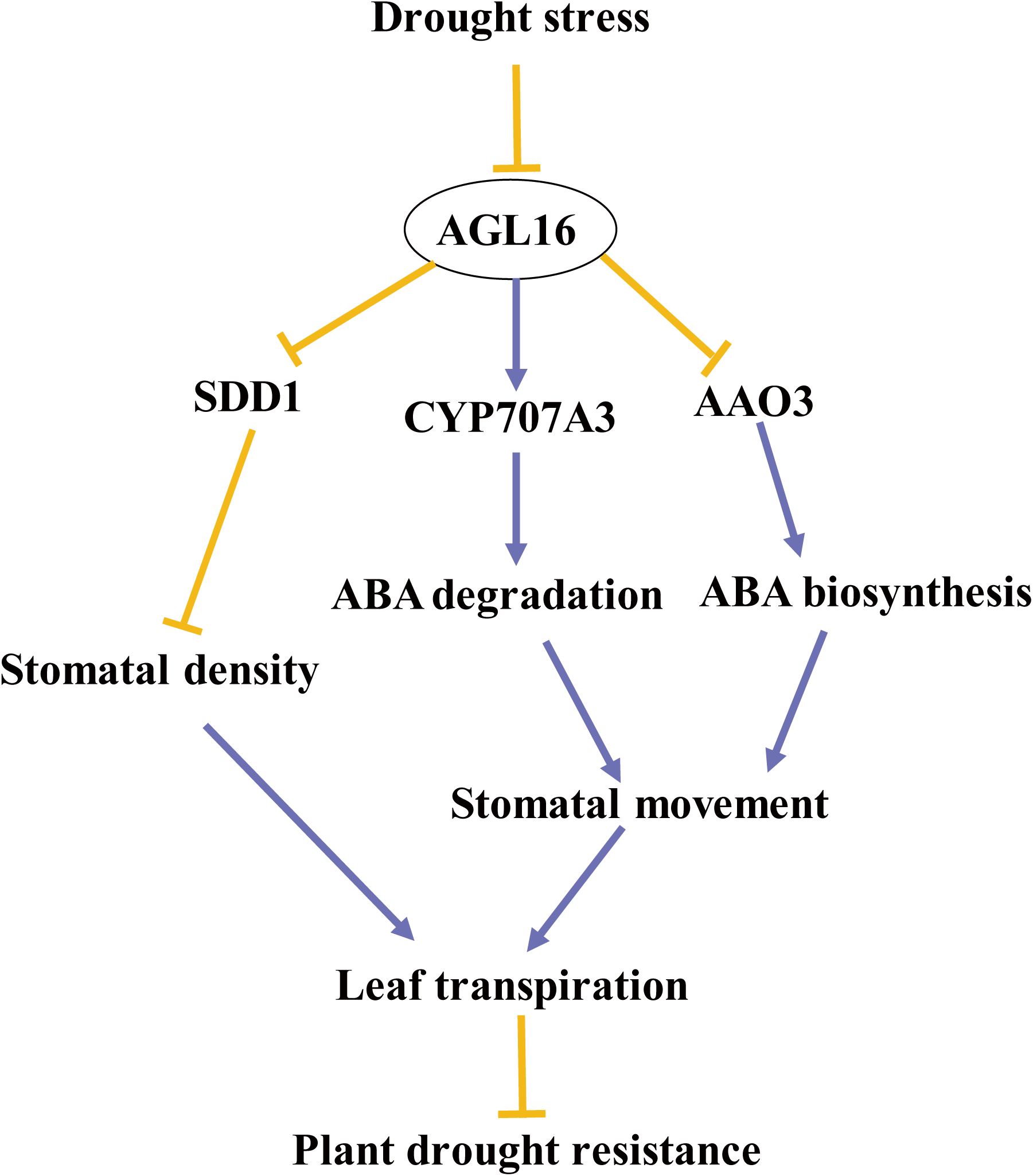
A working model for AGL16 regulating stomatal density and stomatal movement. AGL16 acts as a transcriptional activator to enhance the expression of *CYP707A3* but as a repressor to down regulate the expression of *SDD1* and *AAO3* by directly binding the CArG motifs in their promoter, consequently modulating stomatal density during leaf development and leaf ABA content. In response to drought stress, *AGL16* itself is down regulated, and thus *CYP707A3* is down regulated while *AAO3* is up regulated. As a result, ABA level is increased to promote stomatal closure. On the contrary, when drought stress is diminishing, *AGL16* expression is being increased, leading to decreased ABA levels to favor stomatal opening. Therefore, we propose that AGL16, acting as a negative regulator of drought resistance, is an important modulator balancing between drought resistance and growth.

## MATERIALS AND METHODS

### Plant materials and growth conditions

All *Arabidopsis thaliana* materials used in this study were Columbia ecotype background. A homozygous *AGL16* loss-of-function mutant Salk_104701 was ordered from Arabidopsis Biological Resource Center (ABRC). The *35S:AGL16* overexpression lines, *35S:HA-AGL16*, *AGL16pro: GUS*, *35S:AGL16-GFP* and functional complementation (FC) transgenic plants were obtained by *Agrobacterium* (C58C1) transformation using the Arabidopsis floral-dip method (Clough and Bent, 1998). Arabidopsis seeds were cleaned with 10% bleach for 15-20 minutes and washed by distilled water for 4 times. Seeds were vernalized at 4°C for 2 days, and germinated on MS (Murashige and Skoog) medium vertically for 5 to 7 day, then transplanted into pots to growth. Plants were grown in a suitable environment at 22-24 °C, 60-80% relative humidity, and 16/8h light/dark cycles.

### Leaf RWC analysis

Leaf RWC analysis was conducted as described previously (Yoo et al., 2010). Leaves were detached from plants and weighed to record leaf FW (fresh weight). The leaves were placed in distilled water, fully absorbed and weighed to obtain leaf TW (turgid weight). Then, leaves were dried to reweigh to obtain leaf DW (dry weight). Leaf RWC was calculated as (FW - DW)/(TW - DW)×100.

### Stomatal aperture analysis

Arabidopsis epidermal peels were obtained from the abaxial side of leaves as described in (Zhang et al., 2004). We exposed the sixth leave of Col-0, *agl16* and transgenic plants to open buffer under strong light for 12 h to induce stomatal opening, then epidermal peels were transferred to close buffer with 0, 5, 10 µM ABA for 2 h. Stomata were imaged with HiROX MX5040RZ microscope. Stomatal aperture of different lines were used to analyze in response to ABA (Pei et al., 1997).

### Measurement of ABA Content

Endogenous ABA was extracted from 0.5 g fresh leaves of Col-0, *agl16*, OX and FC transgenic plants under normal or drought stress, and measured by the ABA immunoassay kit as described by (Yang et al., 2001). At least three biological replicates per sample were used.

### ROS activity analysis in guard cells

The ROS were detected in Arabidopsis stomata as described previously (Pei et al., 2000). The lower epidermis of Arabidopsis were incubated in 10 µM 2’,7’-dichlorodihydrofluorescein diacetate (DCFH-DA) solution for 1 hour at 37°C in the dark. Subsequently, lower epidermis were washed by 0.01M PBS buffer which is consisted of NaCl, KCl, Na_2_HPO_4_ and KH_2_PO_4_ twice. ROS-specific fluorescence was observed using ZEISS880 confocal laser scanning microscope with an excitation of 488 nm and an emission of 525 nm condition.

### Histochemical GUS staining

The GUS staining of *AGL16pro: GUS* transgenic plants were performed as described (Jefferson et al., 1987). The mixture was put in 37°C and dark incubator. When GUS staining was finished, the transgenic plants need to destain with 30%, 70%, 100% ethanol and rehydration with 100%, 70%, 30% ethanol. GUS staining solution is composed of X-Gluc, N-N′-Dimethylformamide, 0.1□M sodium phosphate buffer (pH=7.0), 0.5□mM potassium ferrocyanide, 0.5□mM potassium ferricyanide, and 0.1% Triton X-100.

### Subcellular localization of AGL16

*35S:AGL16-GFP* vector was transiently introduced in *Nicotiana benthamiana* epidermal cells by injection as described (Voinnet et al., 2003). Green fluorescence of *35S:*AGL16*-GFP* transgenic leaves were observed by ZEISS880 confocal laser scanning microscope with an excitation of 488 nm and an emission of 525 nm condition. Epidermal peels of transgenic leaves were incubated in 1 µg/mL DAPI solution for 10 min and washed twice with Phosphate buffer saline. Red fluorescence was detected with excitation of 364 nm and an emission of 454 nm condition.

### RT-PCR and quantitative RT-PCR analysis

Total RNA of different tissues was isolated using TRIzol reagent (Invitrogen, Carlsbad, USA). Purified RNA was reversed transcribed by Prime Script RT reagent kit (TaKaRa, Dalian, China). The cDNA was used for the templates of RT-PCR and quantitative RT-PCR. Quantitative RT-PCR was performed by StepOne real-time PCR system using SYBR Premix Ex Taq II kit to detect the expression level of different genes.

### ChIP assay

ChIP assay was performed as previously reported (Cai et al., 2014). 10-day old *35S:HA-AGL16* transgenic seedlings were used in ChIP assay. The seedlings were ground into powder in liquid nitrogen and cross-linked with nuclei isolation buffer. Then chromatin complexes were fragmented into 250-500 bp by sonication. The sonication products were incubated with or without anti-HA antibody (Abmart, Berkeley Heights, USA) and the protein A agarose beads were added into incubation mixtures. The enriched small DNA fragments were eluted from the beads by wash buffer, and the purified DNA was obtained as templates. The enrichment level was detected by PCR and quantitative PCR.

### Transient expression assay

The coding sequence of *AGL16* was cloned into pGreenII 62-SK vector as an effector, the whole promoter of candidate target genes or about 30 bp binding cis-element were respectively cloned into pGreenII 0800-LUC vector as a reporter (Hellens et al., 2005). Arabidopsis mesophyll protoplasts from 4-week wild type leaves were prepared, the effector and reporter were subsequently transfected into mesophyll protoplasts as described by (Yoo et al., 2007). A dual-luciferase reporter assay system was used in transient transfections. The relative activity of firefly LUC and renilla luciferase (REN) was measured, and LUC/REN ratios were calculated. REN was as a control.

### Yeast one-hybrid assay

Yeast one-hybrid assay was carried out as described previously (Mao et al., 2016). The coding sequence of AGL16 was cloned into pAD-GAL4-2.1 (AD vector) and the putative protein binding sites were cloned into pHIS2 (BD vector). These constructs of pAD and pHIS2 plasmids were introduced into Y187 yeast strain by heat shock, and their growth observed on SD/-Trp-Leu medium with or without His.

### Statistical analysis

Statistical analyses were conducted using Student’s t-tests. Values are the mean ± SD and P<0.05 was considered statistically significant (*P<0.05, **P<0.01,).

### Accession numbers

Sequence data from this article can be found in the GenBank/EMBL libraries under the following accession numbers: *AGL16* (At3g57230), *UBQ5* (At3g62250), *CYP707A1* (At4g19230), *CYP707A2* (At2g29090), *CYP707A3* (At5g45340), *CYP707A4* (At3g19270), *ABA1* (At5g67030), *ABA2* (At1g52340), *ABA3* (At1g16540), *NCED3* (At3g14440), *AAO3* (At2g27150), *SDD1* (At1g04110), *ERECTA* (At2g26330), *TMM* (At1g80080), *YODA* (At1g63700), *MPK3* (At3g45640), *FAMA* (At3g24140), *ABI1* (At4g26080), *ABF3* (At4g34000), *RD29B* (At5g52300), *RD22* (At5g25610), *GORK* (At5g37500), *KAT1* (At5g46240), *KAT2* (At4g18290), *AKT1* (At2g26650), *AKT2* (At4g22200), *SLAC1* (At1g12480).

## Supplemental Data

Figure S1. Identification of the loss-off-function mutant *agl16*, AGL16 overexpression lines and functional complementation (FC) transgenic lines

Figure S2. AGL16 affects the leaf temperature by false-color infrared

Figure S3. AGL16 responds to abiotic stress

Figure S4. Expression of ABA signal pathway genes is elevated in *agl16* plants under drought stress.

Figure S5. Expression profiles of outward K^+^ channels and anion channels genes in Col-0, *agl16*, OX plants under drought stress.

## ACKNOWLEDGEMENTS

This study was supported by grants from China Postdoctoral Science Foundation (2019M652200 to P.Z.), NNSFC (31572183 to C.X.) and MOST (2018ZX08009-11B, 2016ZX08005-004-003, 2016ZX08001-003-005 to C.X.). We thank the ABRC for providing the mutant seeds.

## AUTHOR CONTRIBUTIONS

P.Z. and C.X. designed the experiments. P.Z., Z.M., J.Z., Q.L. performed the experiments and data analyses. P.Z. wrote the manuscript. C.X supervised the project and revised the manuscript.

