## supplemental info for "MADS-box factor AGL16 negatively regulates drought resistance via stomatal density and stomatal movement"

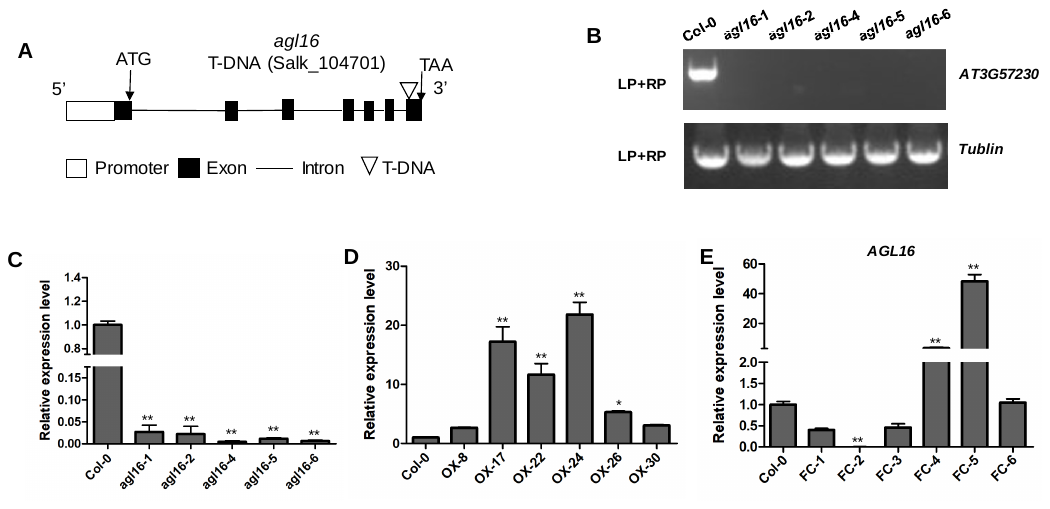

**Figure S1. Identification of the loss-of-function mutant *agl16*, AGL16 overexpression lines and functional complementation (FC) transgenic lines**

(A) Schematic representation of the T-DNA insertion site in Salk_104701. White box represent promoter, black boxes represent exon, black lines represent intron, white triangle represent T-DNA.

(B-C) Identification of genomic and transcript levels of *AGL16* in *agl16* mutant and Col-0. Seeds of Col-0 and *agl16* mutant were germinated and grown on MS medium for 7 days, then DNA and RNA were isolated respectively. PCR and quantitative RT-PCR analysis were performed to detect the genomic (B) and transcript level of *AGL16* (C). Values are mean ± SD (n=3 replicates, Student’s t-test P values, *P<0.05, **P<0.01).

(D) *AGL16* Transcript levels of *35S: AGL16* lines. Seeds of Col-0 and *35S: AGL16* overexpression lines were germinated and grown on MS medium for 7 days, then RNA was isolated and quantitative RT-PCR analyses were performed to detect the transcript level of *AGL16*. Values are mean ± SD (n=3 replicates, Student’s t-test P values, *P<0.05, **P<0.01).
(E) *AGL16* Transcript levels of functional complementation (FC) lines. FC lines were generated by transferring 35S:*AGL16* into *agl16* mutant via Arabidopsis floral-dip method. Seeds of Col-0 and FC lines were germinated and grown on MS medium for 7 days, then RNA was isolated and quantitative RT-PCR analyses were performed to detect the transcript level of *AGL16*. Values are mean ± SD (n=3 replicates, Student’s t-test P values, *P<0.05, **P<0.01).

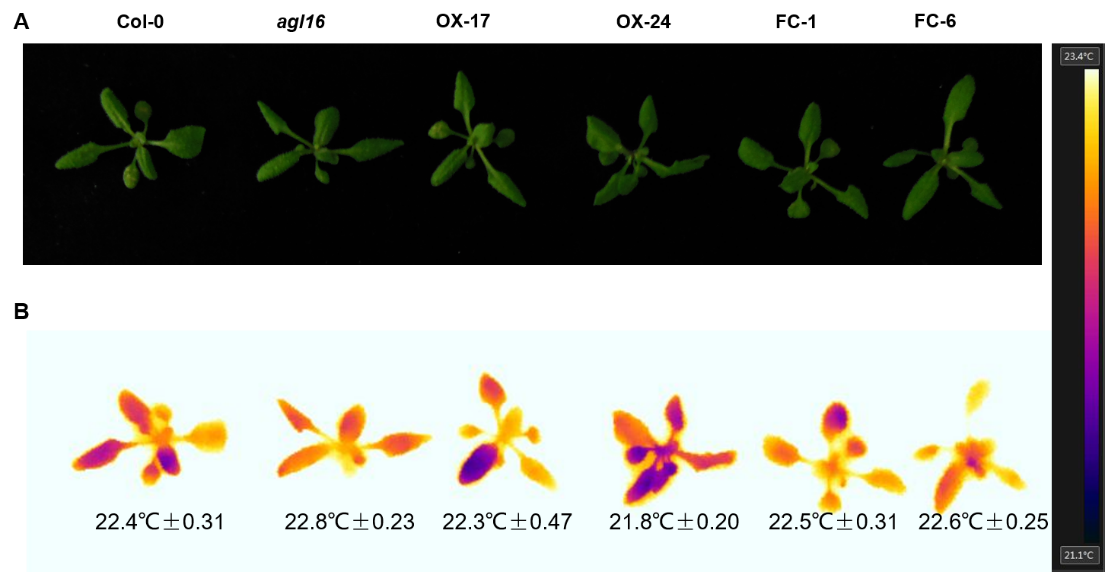

**Figure S2. AGL16 affects leaf temperature by false-color infrared video camera**

The leaf temperature of 3-week-old Col-0, *agl16*, OX, and FC lines were calculated by FLIR Tools software. Photographs were taken under normal condition (A) and false-color infrared images were recorded under the same conditions (B).

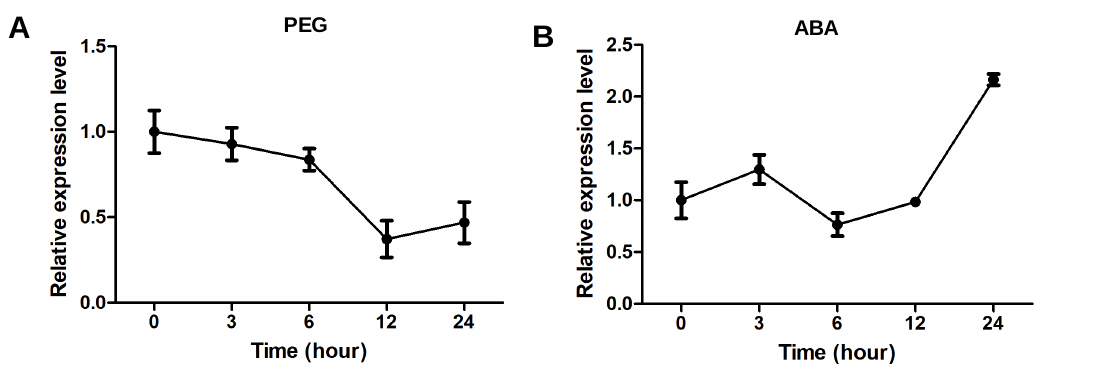

**Figure S3. *AGL16* responds to drought stress and ABA**

2-week-old wild type plants were separately treated with 20% PEG8000 (A) and 10 µM ABA (B) for 0, 3, 6, 12 or 24h. RNA were extracted from shoots to detect the transcript level of *AGL16* by quantitative RT-PCR analysis. Values are mean ± SD (n=3 replicates).

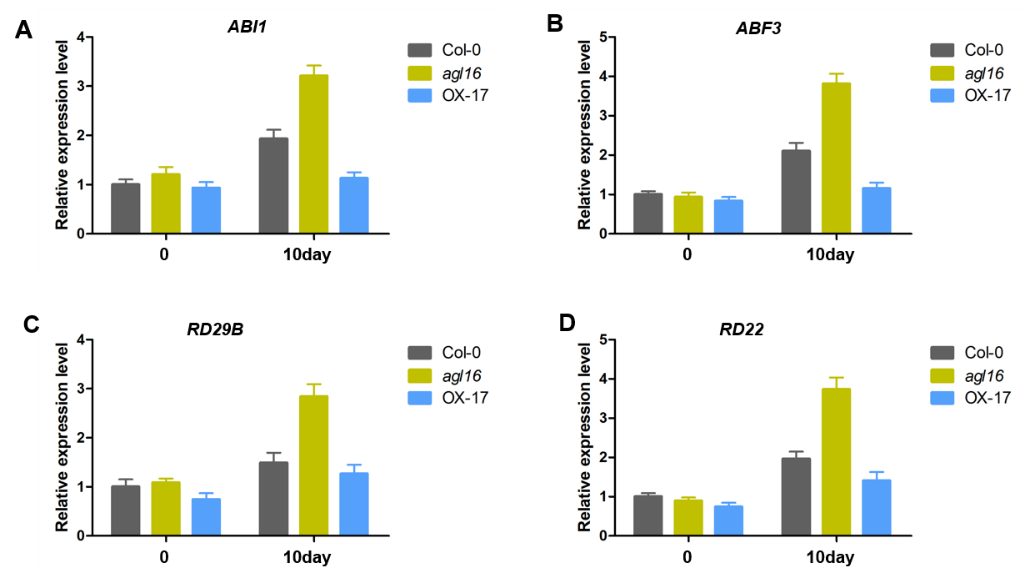

**Figure S4. Expression of ABA-responsive genes is elevated in *agl16* under drought stress** Transcription level of *ABI1*, *ABF3*, *RD29B*, and *RD22* was measured in 4-week-old Col-0, *agl16*, OX plants being withheld water for 0 or 10 days by quantitative RT-PCR analysis. Values are mean ± SD (n=3 replicates).

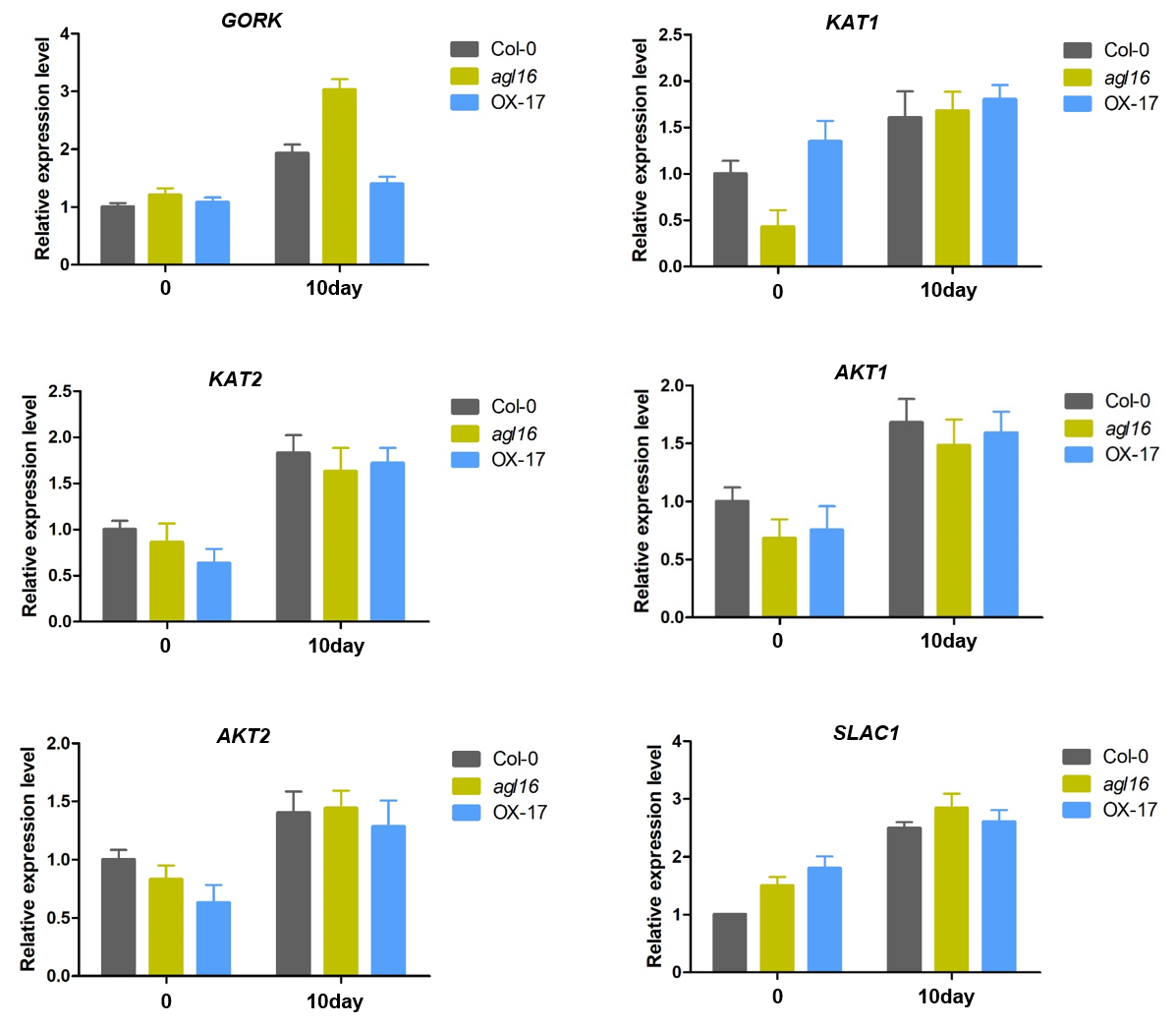

**Figure S5. Expression proﬁles of outward K^+^ channels and anion channels genes in** **Col-0, *agl16*, OX plants under drought stress**

Transcription level of outward K^+^ channels genes *GORK, KAT1, KAT2, AKT1, AKT2* and anion channels gene *SLAC1* were measured in 4-week-old adult Col-0, *agl16*, OX plants under withholding water for 0 or 10days by quantitative RT-PCR analysis. Values are mean ± SD (n=3 replicates).

**Table S1. Primers used in this study**

| Purpose | Primers | Primer sequence |
| --- | --- | --- |
| For OX lines construction | pCB2004-AGL16 LP | GGGGACAAGTTTGTACAAAAAAGCAGGCTATGGGAAGGGGCAAGATCGC |
|  | pCB2004-AGL16 RP | GGGGACCACTTTGTACAAGAAAGCTGGGTTTATGCAATGAAGGAAAAATAG |
| For AGL16pro:GUS construction | GUS-AGL16 LP | GGGGACAAGTTTGTACAAAAAAGCAGGCTAACTATGAACTTGGTAGCTCTTG |
|  | GUS-AGL16 RP | GGGGACCACTTTGTACAAGAAAGCTGGGTTTCTGCTTCTATCACTTTTACAC |
| For 35S:AGL16-GFP construction | GFP-AGL16 LP | GGGGACAAGTTTGTACAAAAAAGCAGGCTATGGGAAGGGGCAAGATCGC |
|  | GFP-AGL16 RP | GGGGACCACTTTGTACAAGAAAGCTGGGTCTGCAATGAAGGAAAAATAGT |
| For HA-tag lines construction | HA-AGL16 LP | GTTTGTACAAAAAAGCAGGCTATGTACCCATACGATGTTCCAGATTACGCTATGGGAAGGGGCAAGATCGC |
|  | HA-AGL16 RP | CTTTGTACAAGAAAGCTGGGTTTATGCAATGAAGGAAAAATAG |
| CYP707A3 promoter ChIP-PCR | CYP707A3 cis1 LP | TGCCCCTTTCCTTGACTTAATCC |
|  | CYP707A3 cis1 RP | CAGTGGAGGAGAGAGAAGGATG |
|  | CYP707A3 cis2 LP | AGTTAAACCTACTTCTATGATTTG |
|  | CYP707A3 cis2 RP | CCACGTTCGCCCTGCATTAC |
|  | CYP707A3 control LP | CAAGATACTCGGACCCAAATCA |
|  | CYP707A3 control RP | GGAAAGGAGGTAATGGGCGG |
| SDD1 promoter ChIP-PCR | SDD1 cis1 LP | CGAGGCGTGTTGAGCGTGCGT |
|  | SDD1 cis1 RP | GTTCACGAGATGTTATGGTTAGG |
|  | SDD1 control LP | ATCTCTTCTGTCGAACATTGGG |
|  | SDD1 control RP | ACAGATCACACTTGAGGGATG |
| AAO3 promoter ChIP-PCR | AAO3 cis2 LP | GAAACTCTTCATCACATCTCCC |
|  | AAO3 cis2 RP | CCAGAGCAGTGACACCAGAAT |
|  | AAO3 control LP | GGTGTTATGAATGAATTGATGGTG |
|  | AAO3 control RP | CAACAACGAACGAGTCTACAGC |
| AGL16 protein for yeast one-hybrid | pAD/AGL16 LP | CGGGATCCATGGGAAGGGGCAAGATCGC |
|  | pAD/AGL16 RP | ACGCGTCGACTTATGCAATGAAGGAAAAATAGTTG |
| CYP707A3 for  yeast-one hybrid | pHIS2/CYP707A3 cis1 LP | CAGCTGGCTCCAAAAAAGGTTCCTTTTA |
|  | pHIS2/CYP707A3 cis1 RP | CGCGTAAAAGGAACCTTTTTTGGAGCCAGCT GAGCT |
|  | pHIS2/CYP707A3 cis2 LP | CTCTAAGCACATATATAGGTTTTGATTA |
|  | pHIS2/CYP707A3 cis2 RP | CGCGTAATCAAAACCTATATATGTGCTTAGA GAGCT |
| SDD1 for  yeast-one hybrid | pHIS2/SDD1cis1 LP | CAAAAAATACATATATTTGGCCAATGGA |
|  | pHIS2/SDD1 cis1 RP | CGCGTCCATTGGCCAAATATATGTATTTTTTGAGCT |
| AAO3 for  yeast-one hybrid | pHIS2/AAO3 cis1 LP | CAATCTAACCTTATAATTGGAAAAATTA |
|  | pHIS2/AAO3 cis1 RP | CGCGTAATTTTTCCAATTATAAGGTTAGATGAGCT |
|  | pHIS2/AAO3 cis2 LP | CATTTCTCTCTTTAATTGGTTAAATCTA |
|  | pHIS2/AAO3 cis2 RP | CGCGTAGATTTAACCAATTAAAGAGAGAAATGAGCT |
| AGL16 protein for transient transactivation assay | pGreenII 62-SK/AGL16 LP | AACTGCAGATGGGAAGGGGCAAGATCGC |
|  | pGreenII 62-SK/AGL16 RP | GCGTCGACTTATGCAATGAAGGAAAAATAGTTG |
| CYP707A3-pro for transient transrepression assay | pGreenII 0800/CYP707A3 LP | GCGTCGACGTCTTTGGTCCTAGATCCCTC |
|  | pGreenII 0800/CYP707A3  RP | AACTGCAGCAGTGGAGGAGAGAGAAGGATG |
|  | pGreenII 0800/CYP707A3 cis1 LP | TCGACAGCTGGCTCCAAAAAAGGTTCCTTTTCTGCA |
|  | pGreenII 0800/CYP707A3  cis1 RP | GAAAAGGAACCTTTTTTGGAGCCAGCTG |
|  | pGreenII 0800/CYP707A3  cis2 LP | TCGACTCTAAGCACATATATAGGTTTTGATTCTGCA |
|  | pGreenII 0800/CYP707A3  cis2 RP | GAATCAAAACCTATATATGTGCTTAGAG |
| SDD1-pro for transient transrepression assay | pGreenII 0800/SDD1 LP | GCGTCGACGAGAGAGTTTTCCACAGGGTTT |
|  | pGreenII 0800/SDD1  RP | AACTGCAGGAAGATGATGAAGCTCGTGAAG |
|  | pGreenII 0800/SDD1 cis1 LP | TCGACAAAAAATACATATATTTGGCCAATGG CTGCA |
|  | pGreenII 0800/SDD1  cis1 RP | GCCATTGGCCAAATATATGTATTTTTTG |
| AAO3-pro for transient transactivation assay | pGreenII 0800/AAO3 LP | GCGTCGACTGCGTTTTATCTTCTTTCCCCA |
|  | pGreenII 0800/AAO3  RP | AACTGCAGTTAACTGCAAACTCCAAATCCAT |
|  | pGreenII 0800/AAO3 cis1 LP | TCGACAATCTAACCTTATAATTGGAAAAATT CTGCA |
|  | pGreenII 0800/AAO3  cis1 RP | GAATTTTTCCAATTATAAGGTTAGATTG |
|  | pGreenII 0800/AAO3 cis2 LP | TCGACATTTCTCTCTTTAATTGGTTAAATCTCTGCA |
|  | pGreenII 0800/AAO3  cis2 RP | GAGATTTAACCAATTAAAGAGAGAAATG |
| For quantitative RT-PCR | Q/Ubiqutin5 LP | AGAAGATCAAGCACAAGCAT |
|  | Q/Ubiqutin5 RP | CAGATCAAGCTTCAACTCCT |
|  | Q/AGL16 LP | GACCTCCACAAGAAAGTAAACC |
|  | Q/AGL16 RP | GCAATGAAGGAAAAATAGTTGAGT |
|  | Q/CYP707A1 LP | AGAGTCTCTAACTTGGGGAGAT |
|  | Q/CYP707A1 RP | TGGATCAAATTTCCCCGGATTA |
|  | Q/CYP707A2 LP | ATTTAAATGGTGGTTGCACTGG |
|  | Q/CYP707A2 RP | GCGAAGAAGGAATTGGGATTTT |
|  | Q/CYP707A3 LP | TGATATATTTTCGGATCCGGGG |
|  | Q/CYP707A3 RP | TTTAGCTAACTCATTGCCTGGA |
|  | Q/CYP707A4 LP | TGCCACTGACACATAAGGTTAT |
|  | Q/CYP707A4 RP | CTCGAATCTAGATGGGTCGAAA |
|  | Q/ABA1 LP | TTGCTGAACAAGTTATGGAAGC |
|  | Q/ABA1 RP | CTGCTGCAGAGTCATTCTACTA |
|  | Q/ABA2 LP | CTTATGTAGTGTGGGAGGTGTT |
|  | Q/ABA2 RP | CTCCTAGTCAAGCCTAGAACAG |
|  | Q/ABA3 LP | AATATGCATTAGCTCAAGGTGC |
|  | Q/ABA3 RP | GCACGATGCTTTACCTTGATAG |
|  | Q/AAO3 LP | TTAGTGTGGCTGAGCTTCATAA |
|  | Q/AAO3 RP | CCAAAGCATCAATAGCATTCGA |
|  | Q/NCED3 LP | GATGAATTTGTTCCAGAGAGCG |
|  | Q/NCED3 RP | AACACTAGGATCAGCCGTTTTA |
|  | Q/SDD1 LP | CAACGGTCGTATTTTCCTATTCA |
|  | Q/SDD1 RP | AGACATGAACTATATTAACTGAAG |
|  | Q/ERECTA LP | TCCTGGTCTTTGCGGTAGTTGG |
|  | Q/ERECTA RP | AGGAGGAGGATTATGCGGTCGG |
|  | Q/TMM LP | ACAAACTGGATTCACGATGACC |
|  | Q/TMM RP | CGTAAGAGCGTTGTGGATCAC |
|  | Q/YODA LP | TCTGTCTGCATCTCCTAGGCGG |
|  | Q/YODA RP | CTGAGTAGCCATATCTCCACCA |
|  | Q/MPK3 LP | CGGCAACTTCCCAACTTCCCA |
|  | Q/MPK3 RP | GAGTGCTATGGCTTCTTGGTAG |
|  | Q/FAMA LP | GGATTGACCCCGTTTATTTCTTG |
|  | Q/FAMA RP | CTTCCTCTTGCTCTTCACCTCC |
|  | Q/ABI1 LP | GTTTGGGATGTAATGACGGATG |
|  | Q/ABI1 RP | ACCACACTTATGTTGTCTTTGC |
|  | Q/ABF3 LP | GATGTGGTTAACCGTTCTCAAC |
|  | Q/ABF3 RP | CAGCTTGCAGTAGATTGTTGTT |
|  | Q/RD29B LP | GAAACCAAAGATGAGTCGACAC |
|  | Q/RD29B RP | TTTTTCGTAAACCGGAGTCAAC |
|  | Q/RD22 LP | GACTTTCGATTTTACCGACGAG |
|  | Q/RD22 RP | CGCTACCGGTTTTACCTTTATG |
|  | Q/GORK LP | CTATCGCGATACTCAGACCTAC |
|  | Q/GORK RP | TATCCACAACAAGTACCTCACC |
|  | Q/KAT1 LP | CACAAAACTCATTTCGGTCACT |
|  | Q/KAT1 RP | GTGGTCGTTAATGTCGTAATGG |
|  | Q/KAT2 LP | TTTGATGGAAAAGGTTGCTCTG |
|  | Q/KAT2 RP | TTTTTCACCTGCAAGTCTTAGC |
|  | Q/AKT1 LP | GCAAGACGTAAAACTAACAACTTC |
|  | Q/AKT1 RP | AACATCTACATCATCAATCTCTGC |
|  | Q/AKT2 LP |  |
|  | Q/AKT2 RP |  |
|  | Q/SLAC1 LP | CCAGTGAAGCTCTTCCTAAAGA |
|  | Q/SLAC1 RP | GTCGAATTGGCCATATATCGTG |
| For PCR | Salk104701C LP | CCATGCATTTTCGGTTTTATG |
|  | Salk104701C RP | CTCTTCTTTTGCCTTTTGCAG |
|  | Tubulin LP | CTTAAGCTCACCACTCCAAGCT |
|  | Tubulin RP | GCACTTCCACTTCGTCTTCTTC |
